# Emotional prosody modulates visual mental imagery

**DOI:** 10.1101/2024.01.10.574936

**Authors:** Weifang Huang, Fengxiang Zhang, Chenghui Zhang, Can Wang, Shiyu Zhang, Yi Pu, Xiang-Zhen Kong

## Abstract

Perceptual stimuli’s emotional properties are vital for human evolution and adaptation. While visual imagery is predominantly regarded as a weak form of perception, the influence of cross-modal emotional features on imagery is still unknown. The present study aims to investigate how emotional prosody modulates imagery quality (i.e., accuracy and clarity) and neural mechanisms using a combination of behavioral tasks and functional magnetic resonance imaging (fMRI). At the behavioral level, our results showed that frustrated conditions induced significant prosody effects on visual mental imagery quality measures, and the effects were particularly pronounced in individuals with lower imagery use tendency. At the neural level, compared with the neutral condition, the emotional prosody conditions (both happy and frustrated) showed stronger activation in various regions including the middle occipital gyrus, supporting the critical role of primary visual system in imagery. Moreover, compared to the frustrated prosody condition, the happy prosody showed stronger activation in the precuneus and anterior cingulate cortex, which are core components of the default mode network. A machine learning prediction analysis with a random forest model identified a significant brain-behavioral correlation between prosody-linked neural activity and individual imagery use tendency. A subsequent Shapley Additive exPlanations (SHAP) analysis further highlighted the primary visual and default mode regions as top contributors to this prediction. Taken together, these results provide new insights for the understanding of how emotional prosody modulates visual mental imagery, considering individual differences, and provide compelling evidence for incorporating emotion as important shaping factor in more general model for imagery.

## Introduction

The environment we live in is characterized with perceptional information of various modalities. Although these different channels of sensation have their own independent representations, cross-modal influence is often unavoidable. For example, a person may be more sensitive to the visual perception of dangerous animals when hearing frightening sounds. Such cross-model interactions have been a focus in the field (e.g., Domínguez-Borràs et al., 2017; Kayser et al., 2008; Zhou et al., 2010).

Emotional features, owing to their pivotal role in evolution and environmental adaptation, stand as crucial attributes of perceptual stimuli (Pacella et al., 2017; Pine et al., 2021). Previous research has mainly focused on exploring the influence of emotional features on perception within the same modality. For instance, positive pictures attract people’s attention more easily compared to neutral pictures, allowing for quicker and more accurate judgments about the image content (Xie & Zhang, 2015). Mental imagery is predominantly considered as a weak form of perception. This ability enables us to perceive the world in mind without directly receiving external stimuli, and plays an important role in a series of cognitive functions, such as navigation, planning, and language comprehension. Although previous studies have researched the influence of emotions on imagery, the experimental designs either made the content of the imagery itself emotional, such as having participants reproduce experiences of disgust in their minds or recreate previously seen emotional pictures or scenes (Köchel et al., 2011; Schienle et al., 2009; Schienle et al., 2008). These experimental designs often have limitations due to the inability to separate the influences of emotion on the encoding and retrieval stages. A very limited studies explored the influence of emotional features on imagery within the same modality, for example, Borst and Kosslyn (2010) presented fearful faces before the imagery task to test the modulation effect of visual emotional stimuli on visual imagery. To our best knowledge, no previous study has focused on the cross-modal impact of emotional stimuli on imagery.

We reason that studying these cross-model interactions is important for understanding the nature of imagery from two aspects. Firstly, it allows for a more realistic simulation of the multisensory interactions that occur in the real world. Imagery is very common and practical in daily life, and stimuli in the real world are inherently complex and multisensory. Thus, studying the cross-modal impact on imagery extends the ecological validity of research beyond the effects of stimuli within the same modality on mental imagery. More importantly, our research aids in gaining a deeper understanding of the nature of imagery, which, akin to perception, possesses a higher degree of flexibility but also exhibits inherent instability (Ishai, 2010; Pearson, 2019). Our study can help us better comprehend why imagery is so variable in real-world settings and the neural mechanisms underlying this variability. This understanding could, in turn, inspire innovative applications of imagery in various practical contexts, such as the treatment of emotional mental disorders (Pearson et al., 2015).

To address this gap, the current study employed a combination of behavioral tasks and functional magnetic resonance imaging (fMRI) using a novel auditorily cued recall task involving verbal paired pictures. Emotional stimuli were introduced through the addition of emotional prosody to the auditory cues. By utilizing prosody as the critical stimulus, we confined the impact of emotional information to the imaging stage and minimized interference from external visual stimuli. Additionally, participants’ tendency to use imagery was measured using the SUIS (Spontaneous Use of Imagery Scale; Kosslyn et al., 1998). At the behavioral level, we focused on participants’ self-reported ratings of imagery quality (i.e., accuracy and clarity) and examined differences between individuals with high and low imagery use tendency. At the neural level, we aimed to identify the brain regions that were modulated by the emotional prosody during the imagining phase and investigate the potential links between the brain activity of these regions and individual differences in imagery behaviors. Note that in the fMRI experiment, only the data from the imagery construction phase, following the completion of hearing the auditory word cue, were included in the analysis of the imagery task. This approach aims to mitigate potential confounding effects on the neural activity of the imagery task due to differences in the prosody carried by the auditory words themselves. Together, these results would provide compelling evidence for the understanding of how emotional prosody modulates visual mental imagery, and its underlying neural correlates and provide new insights for incorporating emotion as one of the most important components in a more general model of mental imagery.

We anticipate that happy and frustrated prosody will exert both similarities and differences in their effects on visual mental imagery. Theoretically, the impact of emotions on visual mental imagery can be attributed to two main factors. Firstly, emotional features may compete for attentional resources, even when they are not directly relevant to the experimental task (Trujillo et al., 2021). Secondly, emotional states or emotional control may also influence cognition (Tyng et al., 2019), and thus influence visual mental imagery construction. Consequently, the presence of emotional prosody, regardless of its specific emotional content, may divert attentional resources that would otherwise be allocated to the imagery task. According to attention capacity theory (Cowan et al., 2005; Tombu & Jolicoeur, 2003), the presence of emotional prosody in the environment is expected to make the imagery task more challenging due to attentional limitations. However, since different prosodies can evoke distinct emotional states, they may engage different neural mechanisms. Taking into consideration prior research showing that positive stimuli often have a more favorable impact on imagery compared to negative stimuli (Cameirão et al., 2016; Wang et al., 2017), we predict that performance under happy prosody will outperform that under frustrated prosody. Additionally, given that individuals with higher SUIS scores are expected to use imagery in their daily life more frequently, thus very possibly getting better training for better resistance to external interference, we anticipate that higher SUIS scores would mitigate the impact of emotional prosody. Consequently, we anticipate that higher SUIS scores would serve to mitigate the impact of emotional prosody on visual mental imagery in the current study.

## Materials and methods

### Participants

In total, eighteen participants (10 females; mean age: 22±3.1 years) were for the norming test, which aimed to creating the stimuli with different emotions and validate prosody effects. Subsequently, we recruited 39 participants (25 females; mean age: 22±2.5 years, SUIS scores: 3.67±0.61) for the behavioral experiment and 31 participants (15 females; mean age: 23±2 years, SUIS scores: 3.44±0.4) for the fMRI experiment. It is important to note that among the 31 participants in the fMRI experiment, five had a self-paced imagery construction phase in their experimental design. To ensure the imagined duration was sufficiently long and consistent with the later uniformly applied fixed duration design, only trials with an imagery construction duration of 2 seconds or more were retained. Additionally, for trials with durations exceeding 4 seconds, only the initial 4 seconds were considered.

The self-reported SUIS scores were collected from each participant after their registration for the behavioral and fMRI experiments. There was no requirement for SUIS scoring during the recruitment for the behavioral experiment. However, for the fMRI experiment, we specifically recruited individuals with lower SUIS scores to maximize the observed effects, based on the findings from the behavioral experiment (see Results).

All participants in our study reported normal or corrected vision, normal or corrected hearing, normal self-reported memory ability, normal self-reported emotion perception, and no history of neurological or psychological illness. Prior to participation, all participants provided informed consent and received moderate financial compensation for their involvement. The study procedures were approved by the Zhejiang University Institutional Review Board.

### Material preparation

#### Auditory cues

Semantic content control. We selected 36 non-animal concrete words from a database of 11,310 simplified Chinese words (Xu et al., 2021) based on their valence and arousal ratings. The arousal ratings in this database range from 0 to 4, with higher values indicating higher levels of activation. Similarly, the valence ratings span from −3 to 3, with larger values representing more positive emotions. To minimize potential entanglement between semantic content from word and emotional information from prosody, we chose words with relatively neutral valence (0.17±0.21) and similar arousal levels (1.61±0.27). Furthermore, three experimenters conducted a thorough check on word familiarity, ensuring that no extreme cases were included.

Auditory parameters control. To control the auditory parameters, each word was recorded by a female native Chinese speaker using either neutral, frustrated, or happy prosody. The recordings were then edited using Audacity ® (Version 3.1.3.) to remove background noise and normalize the peak volume. The duration of each clip was approximately 600ms, with minor variations among prosodies, typically within a range of 200ms. We conducted the norming test for auditory materials to valid their clearness and emotionality (i.e., valence and arousal).

#### Paired pictures

To ensure precise control over the content of mental imagery, we selected pictures from the MultiPic database (Duñabeitia et al., 2018) that shared the same conceptual content as the words. These pictures were grayscale and carefully controlled for complexity.

The well-controlled auditory cues were used in both the behavioral experiment and the fMRI experiment to facilitate participants’ recall of the corresponding memorized pictures, namely to induce mental imagery.

### Experiment procedures

#### Emotional prosody norming test

The norming test was conducted to assess the effectiveness of the auditory cues used in our study, specifically evaluating their clarity, valence, and arousal. Participants completed the test in a quiet room while seated in front of a computer screen with headphones. They were instructed to passively listen to a total of 108 sound clips, including 36 clips each for neutral, frustrated, and happy prosody conditions. For each clip, participants were asked to assess its clarity and provide ratings for arousal and valence.

To evaluate clarity, participants indicated whether they perceived the stimuli as “clear” or “unclear.” Arousal ratings were collected using a five-point scale, where “0” represented a state of being “very calm” and “4” indicated being “very strongly activated” during the imagination phase. Valence ratings were collected using a seven-point scale, with “-3” indicating a “very unpleasant” experience, “0” representing a “neutral” experience, and “+3” indicating a “very pleasant” experience.

#### Behavioral experiment

To establish a standardized source of imagery, participants were instructed to memorize a set of 36 selected pictures along with their corresponding word labels, which were presented in a PDF file.

The order of pictures in the PDF file was randomized, and two versions of the file were created with reversed picture orders to avoid bias. Participants received the PDF file and were instructed to memorize all the pictorial components and basic line orientations of the pictures, emphasizing the need for precise encoding.

To assess participants’ memory accuracy, a recognition test was administered using the online platform Palvlovia (https://pavlovia.org/). Participants had to determine which picture out of a pair matched the one they were asked to remember. Only participants with an accuracy rate higher than 95% in this recognition test were eligible to proceed to the main experiment, which focused on visually imaging the picture.

In the main experiment, participants completed a total of 36 trials, with 12 trials for each of the three prosody conditions (neutral, frustrated, and happy). The trials with the same prosody condition were grouped together in blocks to minimize any carry-over effects. The order of the prosody blocks varied to counterbalance the presentation order, with the neutral block always appearing first to establish a baseline for recognizing emotional prosody later. Trial order within each block was randomized. Short breaks of 6 seconds were provided after every 4 trials within a block, and a longer break of 20 seconds was given between each block to mitigate fatigue.

Before running the experiment, the participants were asked to listen to instructions and to finish practice first. To minimize participants’ confusions when perceiving emotional prosody since some emotional prosodies had similar acoustic parameters but were referred to as different emotion type, for example, “thrilled” and “happy,” all participants had prior knowledge, that the auditory cues they heard might contain different prosodies, namely neutral, frustrated, and happy.

The participants were tested using PsychoPy 2022.2.1 (https://www.psychopy.org/) in a quiet room sitting approximately 60cm from the computer screen with headphones on. Each trial began with the presentation of an auditory cue for an average duration of 600ms. Participants were instructed to construct the corresponding visual mental imagery as vividly as possible while keeping their eyes open and focusing on the screen. They were asked to rate the clarity of their constructed mental imagery on an eleven-point scale, where “0” represented “very vague” and “10” represented “very clear”. Following their response, a key feature judgment question related to the constructed imagery was presented on the screen. For example “杯子中是否有水? (Translated in English ‘Is there water in the glass?’)”. An equal number of “yes” and “no” responses were required across the 36 judgments.

After the key feature judgment, the picture that participants were asked to memorize prior was displayed for 1000ms as a correction. Subsequently, participants were shown two ratings to assess the similarity between their constructed mental imagery and the corrected picture. They were asked to rate the similarity in terms of “component” and “line orientation,” where “0” represented “very dissimilar” and “10” represented “very similar,” which served as more operative measures of imagery accuracy. Arousal and valence ratings, using the same scale as the norming test, were collected at the end of each trial to evaluate participants’ emotional feelings during the imagery phase. For a detailed overview of the experimental design, refer to Fig.1(a).

**Fig. 1.**
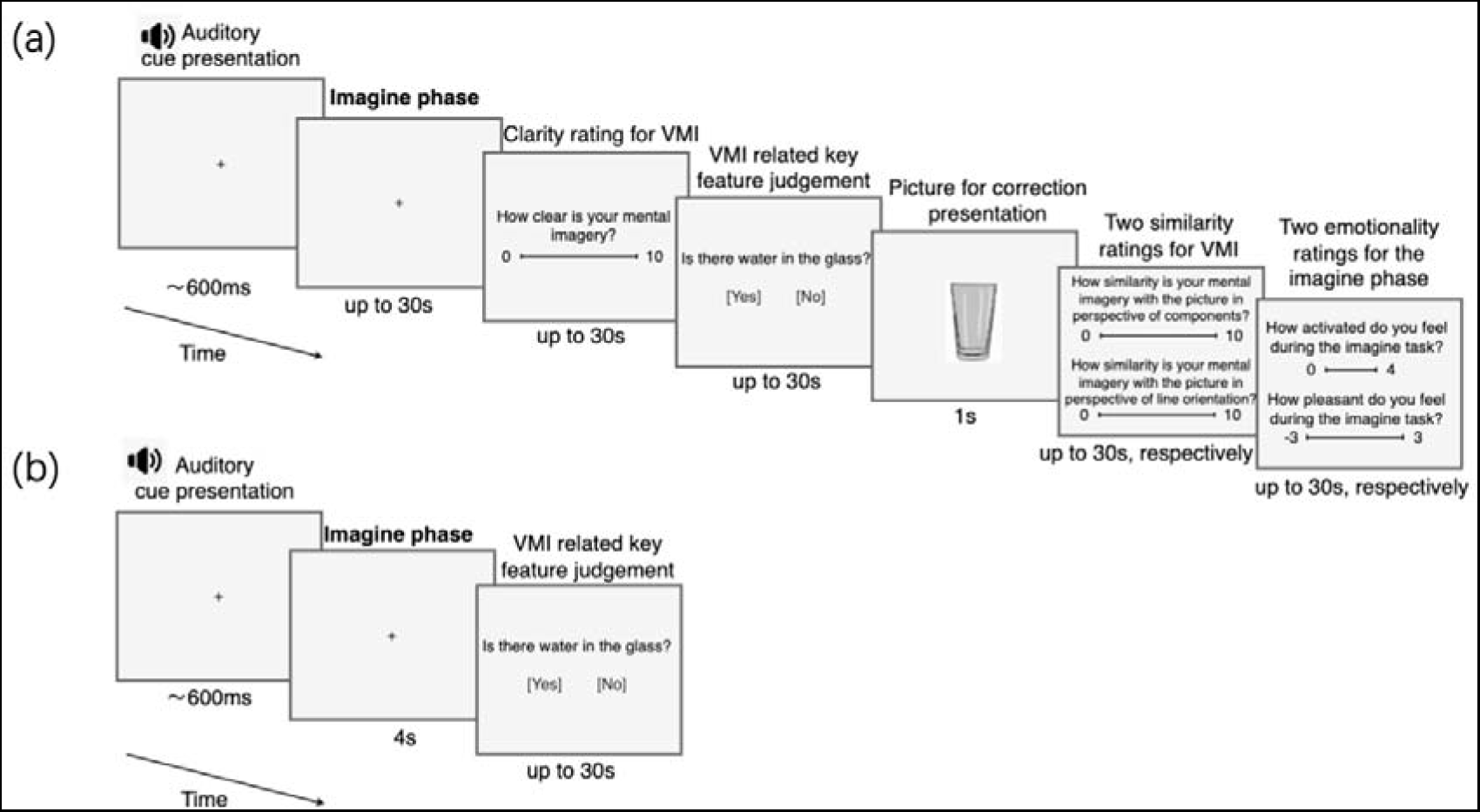
One-trial procedure employed in both the behavioral and fMRI experiments. In contrast with the version of the behavioral experiment (a), the version of the fMRI experiment (b) was simplified, to accommodate the constraints of conducting tasks in the fMRI setting and reduce confounding effects caused by task-related motion and complex cognitive activities. The imagination period in the fMRI experiment lasts for fixed 4 seconds, based on the results of the behavioral study, which confirmed its adequacy for imagination.

Previous research has shown that individuals have robust metacognition for visual mental imagery (Pearson et al., 2011), which implies that people, particularly those with normal visual mental imagery abilities, possess the capacity to recognize their mental imagery and could make valid judgments concerning quality-related aspects. Although using an 11-point scale (0-10 rating) to assess imagery quality is less commonly employed, we chose this scale for two compelling reasons. Firstly, the 11-point scale is considered to be as valid as other commonly used scales, such as 5-point or 7-point scales (Lawless et al., 2010; Leung, 2011). Secondly, our experimental setup involved imagery tasks with clearly defined “standard answers.” As a result, the 0-10 rating scale more accurately reflected the natural thought processes of participants, where a rating of 0 indicated no similarity at all, and a rating of 10 represented an exact match.

The duration of self-paced operations was limited to a maximum of 30 seconds. Apart from predetermined rest intervals, the interval between trials was approximately 1.5 seconds. A fixation cross was consistently presented on the screen, except during the rating and judgment trials. Participants were instructed to maintain their gaze on the fixation cross, except during the imagery task, where flexible eye movements within the screen were allowed. This flexibility in eye movements during imagery was supported by previous research indicating that restricting eye movements can hinder the quality of imagery (Bourlon et al., 2011; Laeng & Teodorescu, 2002; Mast, 2002).

#### fMRI experiment

The fMRI stimuli were identical to those used in the behavioral experiment (i.e., the auditory cue, as well as the to be memorized pictures). Practice trials were finished before entering the scanner. Prior to the main task, participants underwent a 40-second period of rest in the scanner, during which they were instructed to clear their minds and fixate on a cross displayed on the screen. The data collected during this stage served as the baseline measurement. Subsequently, a message appeared on the screen for 2000ms, signaling the start of the main task.

During each trial, participants engaged in a fixed 4-second period of mental imagery immediately after hearing the auditory cue. Following the imagery session, participants promptly performed key feature judgments to assess trial effectiveness. This simplified procedure aimed to minimize the influence of brain activity resulting from rating tasks and any potential movement artifacts caused by task responding. For specific details of the experimental design, refer to Fig.1(b). The self-paced timing for judgments and the interval between trials remained consistent with the procedures used in the behavioral experiment.

MRI scans were conducted using a 3T Siemens MRI scanner with a 64-channel head coil (Siemens Medical Systems, Erlangen, Germany) at the Institute of Translational Medicine, Zhejiang University. Participants lay in the scanner while holding a button box in their right hand. Foam pads were used to minimize head motion. Whole-brain fMRI data were acquired using an Echo Planar Imaging (EPI) sequence, consisting of 52 axial slices of 1.6mm thickness with no gap. The acquisition parameters were as follows: repetition time (TR) = 1000ms, echo time (TE)=34ms, flip angle (FA)=50, field of view (FOV)=260mm, matrix=92x92, and interleaved acquisition. After the functional MRI scan, whole-brain structural scans were obtained using an MP-RAGE anatomical sequence, comprising 208 axial slices of 1mm thickness. The parameters for the structural scans were as follows: TR = 2300ms, TE= 2.32ms, FA=8, FOV= 240mm, matrix=256x256, and interleaved acquisition.

#### Analysis for the behavioral data

In the norming test, we calculated the average percentage of “clear” responses for each type of prosody across participants. The results show that the auditory stimuli achieved a high level of clarity, with the average percentage of “clear” responses exceeding 95% for each prosody condition. To ensure the validity of the auditory materials, we checked for stimuli rated as unclear by more than two participants. No stimuli were excluded based on this criterion. To account for the non-normal distribution of the data (see Results), we used the Wilcoxon test for subsequent statistical analysis, which was performed using Python 3.9.5.

In the behavioral experiment, we collected emotionality ratings using a similar approach as in the norming test. The analysis was conducted using the Wilcoxon test, and Python 3.9.5 was utilized for data analysis. Initially, we analyzed the results related to valence and arousal for all participants, aiming to replicate the results from the norming test. Subsequently, we categorized participants into two groups based on the median SUIS score (3.68). The lower tendency group consisted of 21 participants (15 females; age: 21±2.5 years; SUIS scores: 3.21±0.43), while the higher tendency group included 18 participants (11 females; age: 22±2.5 years; SUIS scores: 4.20±0.27). The purpose was to investigate whether the emotionality resulting from emotional prosody would be influenced by their imagery use tendency.

Additionally, we analyzed the results of the imagery quality ratings in the behavioral experiment using paired t-tests for all participants, as well as separately for the higher and lower tendency groups. Categorizing participants into these groups allowed us to directly examine whether the influence of emotional prosody on imagery quality would be moderated by their imagery use tendency.

To ensure the trials indeed reflecting participants’ imagination of the corresponding picture, only trials with correct key feature judgments were included. Among all participants, the accuracy rates for the neutral, happy, and frustrated prosody conditions were 87%, 85%, and 83%, respectively. For participants with lower SUIS scores, the accuracy rates for all three prosody conditions were 85%. For participants with higher SUIS scores, the accuracy rates for the neutral, frustrated, and happy prosody conditions were 89%, 84%, and 81%, respectively.

### Analysis for the imaging data

#### Preprocessing

Image preprocessing was performed using Statistical Parametric Mapping 12 (SPM12, Wellcome Department of Imaging Neuroscience, London, UK: https://www.fil.ion.ucl.ac.uk/spm/software/spm12/) in Matlab R2017a. The first 10 volumes were discarded to allow the signal to reach a steady state. The remaining volumes underwent the following preprocessing steps: (1) slice timing correction, (2) realignment for motion correction to the mean image, (3) coregistration between the structural and functional data, (4) segmentation of the structural image using default tissue probability maps as priors with East Asian brains as affine regularization, and with forward deformation fields, (5) normalization of the structural and functional images to the standard Montreal Neurological Institute (MNI) space, and (6) smoothing using a 5x5x5mm full width half maximum (FWHM) Gaussian kernel.

#### Whole-brain analysis and contrasts of interest

In line with the behavioral experiment, only trials in which the key feature judgment was answered correctly and baseline data were included in the subsequent analysis. The judgment accuracy for the neutral, frustrated, and happy prosody conditions was 86%, 89%, and 87%, respectively.

At the single-subject level, a fixed general linear model was applied in SPM12 to generate task activation maps for three prosody condition contrasts: frustrated vs. neutral, happy vs. neutral, and happy vs. frustrated. For the prosody condition, data from the imagery task were restricted to the 4-second duration of imagery construction, aiming to minimize the potential confounding effects of variations in prosody within the auditory word cues. In the analyses, trials where participants provided incorrect key feature judgments were deemed unsuccessful and consequently excluded from the analysis. Six rigid-body head motion parameters were included as multiple regressors to address potential physiological artifacts.

Group-level analysis was performed using one-sample t-tests in SPM12. All contrasts were generated with an uncorrected threshold of *p*<0.001 at the voxel level and corrected for false discovery rate (FDR) at the cluster level (*p*<0.05), except for the happy vs. frustrated prosody contrast, which had an uncorrected threshold of *p*<0.005 at the voxel level with FDR of *p*<0.05 at the cluster level.

In addition, we also analyzed contrast maps for each prosody condition compared to the baseline to assess the visual imagery effects.

#### Region-of-interest analyses

To further understand the specific activities resulted from the prosody contrasts, we created 12 ROIs from clusters which survived the multiple testing correction (see Tab.1). Specifically, there were 6 ROIs for contrasting frustrated vs. neutral prosody condition, 3 ROIs for contrasting happy vs. neutral prosody condition, and 3 ROIs for contrasting happy vs. frustrated prosody condition.

Average beta values within these ROIs were then extracted, from happy vs. baseline, frustrated vs. baseline, and neutral vs. baseline conditions resulted SPM files, for each participant. Bar plots visually present the comparison of activation levels of each prosody condition (in relative to baseline) in the prosody contrast resulted ROIs.

#### Random-forest model construction for SUIS scores prediction

To investigate whether the brain activity observed in the prosody contrast was modulated by participants’ imagery use tendency, we extracted the mean beta values for each ROI, which was resulted from the corresponding prosody contrast (frustrated vs. neutral, happy vs. neutral, frustrated vs. happy). The beta values for each ROI were used as features to construct a predictive model for SUIS scores based on a random forest model. Leave-one-out cross-validation was employed to assess the predictive performance of the model. The SHAP (SHapley Additive exPlanations) model (Lundberg & Lee, 2017) was then utilized to demonstrate each feature’s importance in predicting SUIS scores.

## Results

### Validation of emotional effects of the prosody conditions

We first investigated the emotional effects of the prosody conditions established in the present study. In the norming test, significant differences were observed in both the arousal and valance dimensions among the three prosody conditions. Specifically, in terms of arousal, the happy prosody condition received the highest scores, followed by the frustrated prosody condition, and the neutral prosody condition (happy vs. frustrated: *p=*1.88e-5; frustrated vs. neutral: *p=3*.72e-30; happy vs. neutral: *p*=1.08e-75). Similarly, in terms of valence, significant differences were found among the different prosody conditions. The happy prosody condition received the highest scores, followed by the neutral prosody condition, and the frustrated prosody condition (happy vs. neutral: *p*= 4.14e-31; neutral vs. frustrated: *p*=9.45e-95; happy vs. frustrated: *p*=5.07e-97) (Fig. 2a).

**Fig. 2.**
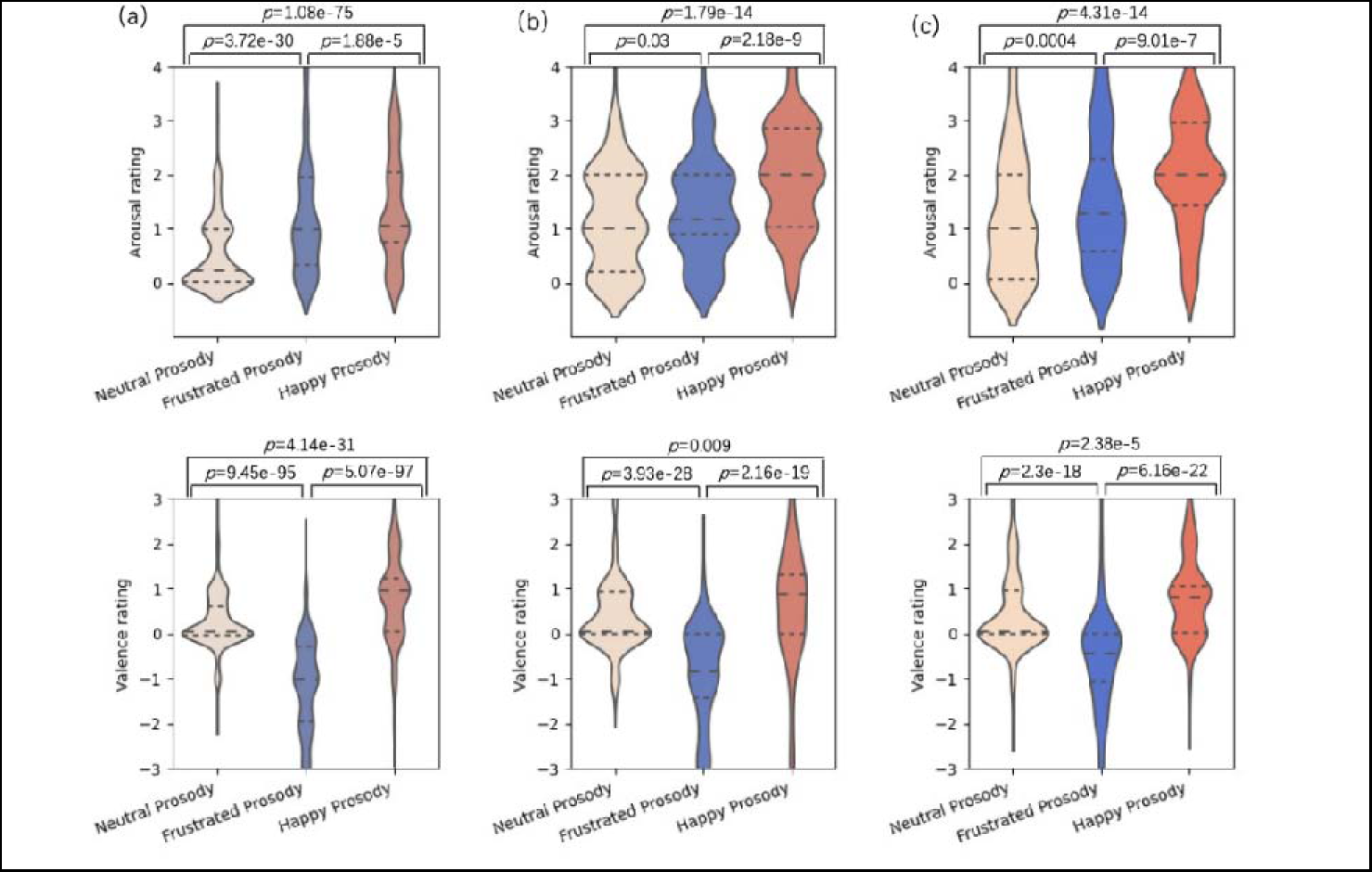
Emotionality rating results for the norming test and the behavioral experiment. (a) Norming test results (N=18), depicting arousal and valence ratings in violin plots. Quartiles are represented by dashed lines. Density estimation is applied beyond the original rating scales. Only trials in which participants reported clear perception were included in the analysis. Wilcoxon non-parametric tests were conducted to examine group differences between prosody conditions, with significant differences denoted by directly labeled p-values. (b) Emotionality rating results in the behavioral experiment, restricted to participants with lower SUIS scores (N=21). Statistical analysis and annotation of significant differences followed the same approach as in (a). (c) Emotionality rating results in the behavioral experiment, limited to participants with higher SUIS scores (N=18). Statistical analysis and annotation of significant differences followed the same approach as in (a). Results plots for all participants are provided in the supplementary materials (see Supp Fig.1).

Similar results were observed in the imagery task. Specifically, in the arousal rating, the happy prosody condition obtained the highest scores, then the frustrated prosody, and the neutral prosody condition (for happy vs. frustrated, p=1.63e-14, Wilcoxon; for frustrated vs. neutral, p=0.0001, Wilcoxon; for happy vs. neutral, p=8.58e-27, Wilcoxon). In the valence rating, the happy prosody condition obtained the highest scores, then the neutral prosody, and the frustrated prosody condition (for happy vs. neutral, p=3.7e-6, Wilcoxon; for neutral vs. frustrated, p=3.53e-44, Wilcoxon; for happy vs. frustrated, p=7.48e-39, Wilcoxon). These results served as validation for our prosody conditions. For specific plots see Fig. S1.

Moreover, when dividing the participants into high and low groups based on the SUIS scores, we found consistent patterns (Fig. 2b, 2c). These results indicated that the prosody effects remained resistant to individuals with different imagery use tendency, which lay the groundwork for further investigation of how emotional prosody modulates visual mental imagery.

### Behavioral effects of prosody on visual mental imagery

Next, we investigated whether prosody modulates imagery quality. Imagery quality was measured by clarity rating, similarity rating for component, and similarity rating for line orientation (see Materials and Methods). The results show that, while there were no significant differences between the happy prosody and the neutral prosody condition (uncorrected *p*s >0.05), the frustrated prosody condition had significantly or marginally significantly lower rating scores than the neutral prosody (clarity rating: *t*(38)=-2, *p*=0.05, *d*=-0.25; similarity rating for line orientation: *t*(38)=-1.97, *p*=0.06, *d*=-0.33). Furthermore, the frustrated prosody condition had significantly lower rating scores than the happy prosody condition in all three ratings (clarity rating: *t*(38)=-3.12, *p*=0.003, *d*=-0.36; similarity rating for component: (*t*(38)=-2.28, *p*=0.03, *d*=-0.35; similarity rating for line orientation: *t*(38)=-2.76, *p*=0.008, *d*=-0.44). These results indicate that while frustrated prosody has a detrimental impact on imagery quality, happy prosody does not cause any impairment to imagery quality.

We further investigated how individual’s imagery use tendency modulates the prosody effects. We found that the observed prosody effects on imagery were mainly contributed by individuals with lower imagery ability. Specifically, in the lower SUIS group, the happy prosody resulted in better imagery quality ratings compared with the frustrated, i.e., clarity rating: *t*(20)=3.99, *p*=0.0007, *d*=0.53; similarity rating for component: (*t*(20)=2.5, *p*=0.02, *d*=0.54; similarity rating for line orientation: *t*(20)=3.13, *p*=0.005, *d*=0.69. In contrast, no significant differences were found in the higher SUIS group.

These results suggest that emotional prosody differences (i.e., happy vs. frustrated) modulate imagery quality, but the effects are linked to individual’s imagery use tendency. Individuals with lower use tendency would be more susceptible to the impact of emotional prosody.

### Neural correlates of the prosody effects on mental imagery

After that, we investigated the neural correlates underlying the observed effects in behaviors. In the fMRI experiment, we selectively recruited a group of participants with lower imagery ability indexed by the SUIS scores according to the behavioral findings of the contrast between the happy and frustrated prosody. The results in this section are based on the performance from 31 participants. For the analysis and results specific to the 26 participants who utilized a fixed duration of 4 seconds for imagery construction, see the supplementary materials.

The visual imagery conditions showed distributed activation across the brain compared with the baseline, and the activation pattern was similar among the three conditions (Fig. Fig. S2). These were in line with previous findings (Pearson, 2019). Moreover, the fMRI contrasts of happy-neutral and frustrated-neutral showed significant activation in multiple regions. Specifically, the frustrated prosody condition showed stronger activation in regions of the left lingual gyrus (MNI coordinates [−14, 97, −12], *t*=5.61, *p*<0.001), bilateral superior temporal gyrus (left: MNI coordinates [−64, −24, 8], *t*=5.2, *p*=0.000; right: MNI coordinates [58, −14, −5], *t*=5.08, *p*=0.000), left inferior frontal gyrus, pars triangularis (MNI coordinates [−42, 16, 28], *t*=5.17, *p*<0.001), right inferior frontal gyrus, pars opercularis (MNI coordinates [53, 23, 32], *t*=4.23, *p*=0.014), as well as left precentral regions (MNI coordinates [−40, 3, 58], *t*=4.05, *p*=0.049) (Table 1, Fig. 4b). Similarly, the happy prosody condition showed stronger activation in regions of the right lingual gyrus (MNI coordinates [6, −87, −5], *t*=6.14, *p*=0.000), right middle occipital gyrus (MNI coordinates [23, −97, 5], *t*=4.54, *p*=0.012), and left superior temporal gyrus (MNI coordinates [−64, −22, 5], *t*=6.1, *p*<0.001) (Table 1, Fig. 4b).

**Fig. 3.**
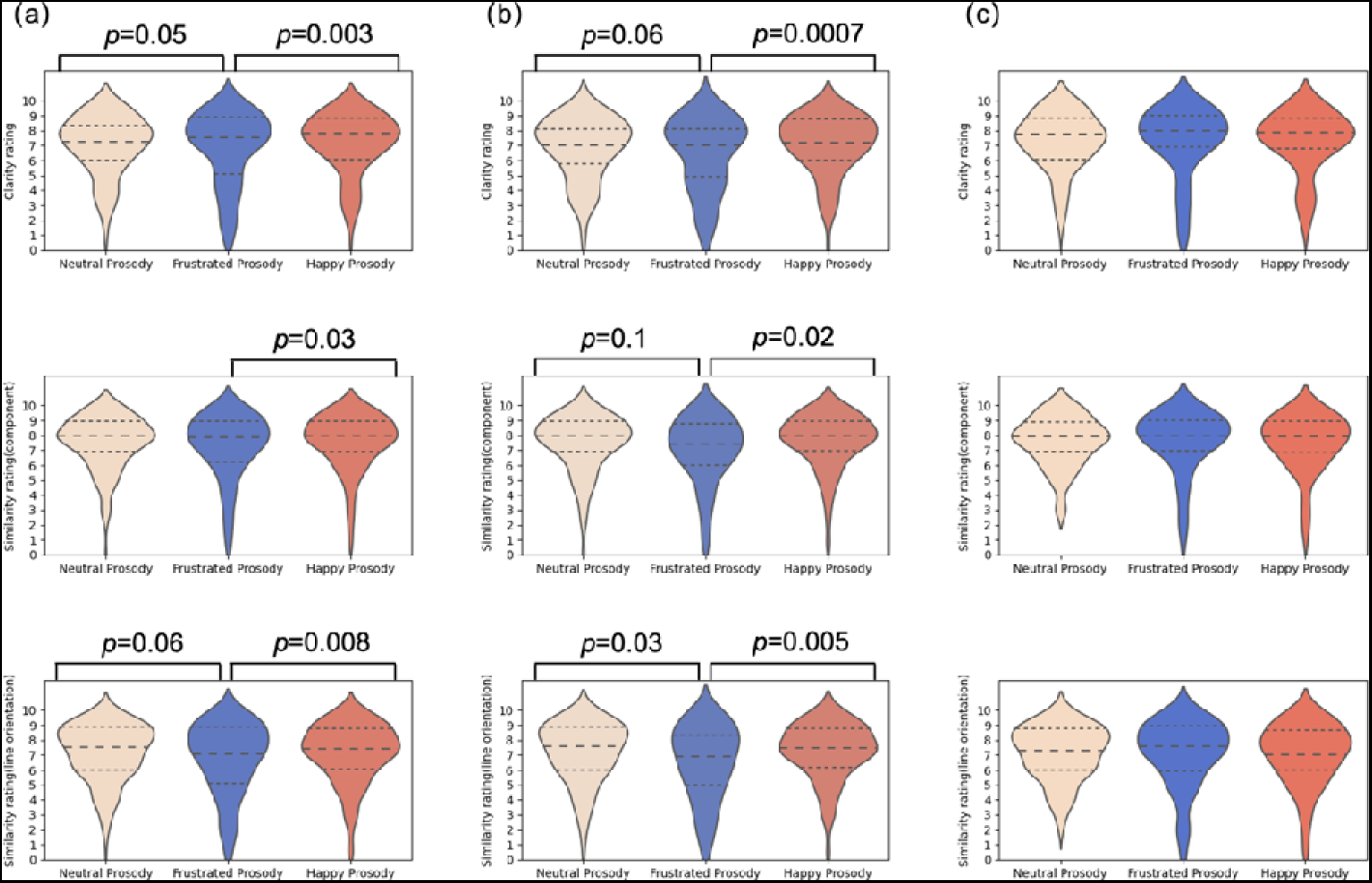
Imagery quality rating results in the behavioral experiment. (a) Rating results for all participants (N=39), including only trials with correct key feature judgments. Violin plots depict the distribution of ratings, with quartiles represented by dashed lines. Density estimation is applied beyond the original rating scales. To enhance the operationality of imagery quality, clarity and accuracy dimensions were assessed. Given fixed imagery sources (i.e., memorized pictures), accuracy is equivalent to similarity with the picture, further subdivided into similarity for component and similarity for line orientation to improve operability for participants. Paired t-tests were conducted for each rating dimension between prosody conditions, with significant or marginally significant differences denoted by labeled *p*-values. (b) Rating results for individuals with lower SUIS scores (N=21). The participants were divided into high-scoring and low-scoring groups based on the median score of the SUIS data from 39 participants. Data analysis and annotation followed the same approach as for all participants. (c) Rating results for individuals with higher SUIS scores (N=18). Data analysis and annotation followed the same approach as for all participants.

**Fig. 4.**
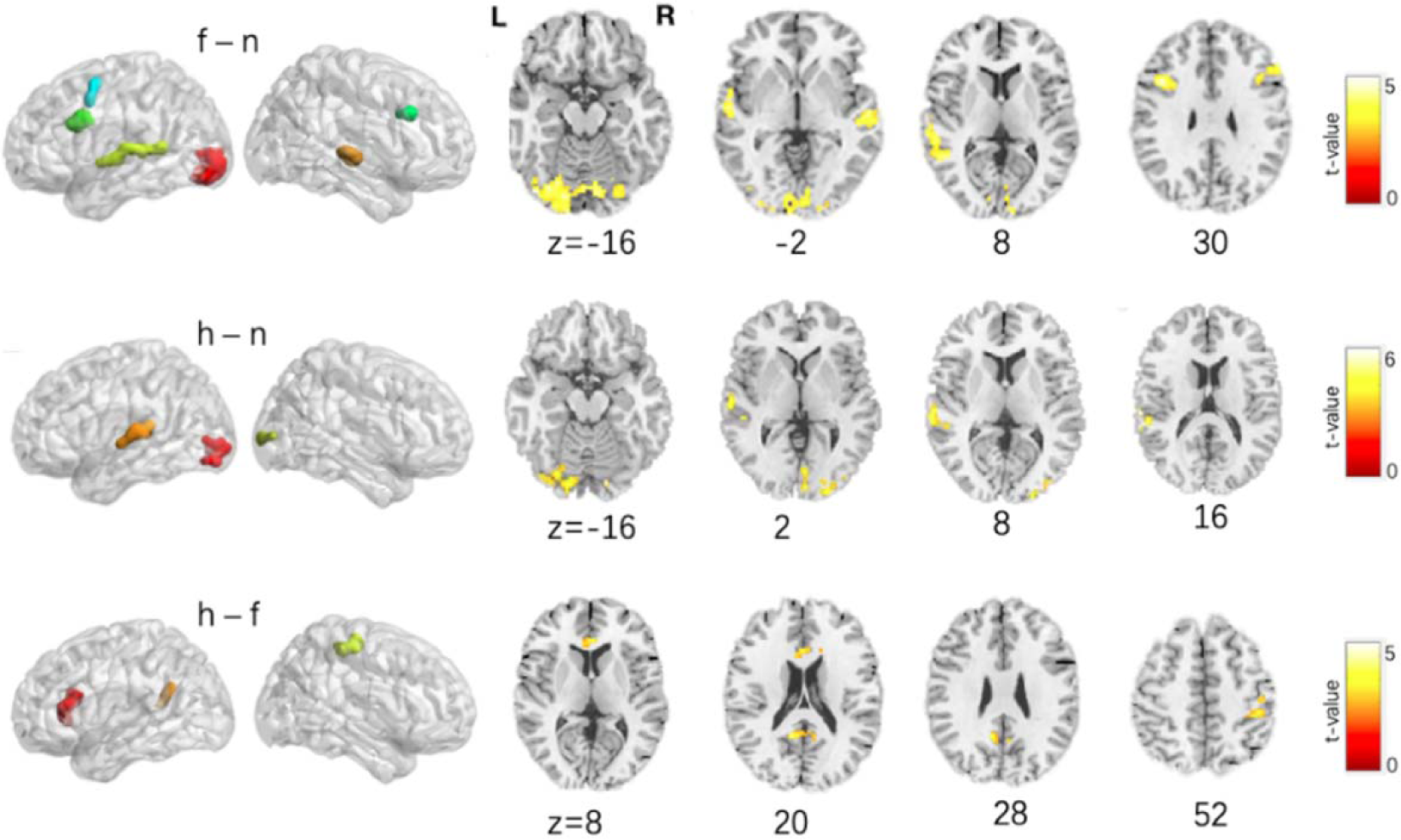
fMRI results for the prosody effects on visual imagery. Activations for prosody condition contrasts, including frustrated(f)-neutral(n); happy(h)-neutral(n), and happy(h)-frustrated(f). All clusters, except for the h-f contrast, were generated using an uncorrected p-value threshold of <0.001 and corrected for false discovery rate (FDR) at the cluster level (*p* < 0.05). Clusters for the h-f contrast were generated using an uncorrected *p*-value threshold of <0.005 and corrected for FDR at the cluster level (*p* < 0.05).

**Tab. 1.** Significant clusters for prosody condition contrasts. Abbreviations “n,” “f,” and “h” represent the neutral, frustrated, and happy prosody conditions, respectively, in the cluster results. All clusters, except for h-f, were generated using an uncorrected *p*-value threshold of <0.001 and corrected for false discovery rate (FDR) at the cluster level (*p* < 0.05). Clusters for the h-f contrast were generated using an uncorrected *p*-value threshold of <0.005 and corrected for FDR at the cluster level (*p* < 0.05).

| Contrast | Cluster | Cluster size<br>(voxels) | Peak MNI<br>coordinates | $t$-value | Cluster $p$-<br>value (FDR) |
| --- | --- | --- | --- | --- | --- |
| f-n | Lingual_L | 957 | -14 -97 -12 | 5.61 | $<0.001$ |
| | Temporal_Sup_L | 215 | -64 -24 8 | 5.20 | $<0.001$ |
| | Frontal_Inf_Tri_L | 141 | -42 16 28 | 5.17 | $<0.001$ |
| | Temporal_Sup_R | 128 | 58 -14 -5 | 5.08 | $<0.001$ |
|  | Frontal_Inf_Oper_R | 51 | 53 23 32 | 4.23 | 0.014 |
|  | Precentral_L | 36 | -40 3 58 | 4.05 | 0.049 |
| h-n | Lingual_R | 285 | 6 -87 -5 | 6.14 | $<0.001$ |
| | Temporal_Sup_L | 119 | -64 -22 5 | 6.10 | $<0.001$ |
|  | Occipital_Mid_R | 66 | 23 -97 5 | 4.54 | 0.012 |
| h-f | ACC_sup_L | 148 | -2 23 15 | 5.26 | 0.014 |
|  | Precuneus_L | 97 | -12 -52 22 | 4.84 | 0.045 |
|  | Precentral_R | 116 | 28 -20 60 | 4.02 | 0.028 |

In terms of the happy-frustrated contrast, significant activations were found in regions of the left anterior cingulate cortex (MNI coordinates [−2, 23, 15], *t*=5.26, *p*=0.014), left precuneus (MNI coordinates [−12, −52, 22], *t*=4.84, *p*=0.045), and right precentral gyrus (MNI coordinates [−12, −52, 22.5], *t*=4.84, *p*= 0.028) (Table 1, Fig. 4b). Interestingly, these regions cover the core components of the default mode network (e.g., Raichle 2015).

### Prosody-linked neural activity predicts individual differences in imagery use tendency

As observed above, individuals with different imagery ability showed variability in prosody effects on quality of mental imagery. Thus, we hypothesized that variability in prosody-linked neural activity during imagery could reflect individual differences in imagery use tendency. We constructed a machine learning prediction model to predict individual’s SUIS score based on beta values of the regions revealed in the fMRI experiment (see Materials and Methods). In total, 12 regions were included in the modelling. The predicted scores showed significant correlation with the actual scores (*r* = 0.40, *p* = 0.025). Moreover, the SHAP analysis showed various contribution scores of different regions. The results unveiled that the activations in the right middle occipital region during the frustrated-neutral contrast and the happy-neutral contrast had a significant negative contribution to the prediction of SUIS scores. That is, higher activation in the middle occipital region indicates a decreased visual imagery use tendency and lower activation indicates an increased tendency. Conversely, the anterior cingulate cortex in the happy-frustrated contrast exhibited a significant positive contribution, while the contribution of the precuneus was relatively mixed (see Fig.6). These brain-behavioral correlation results further supported the functional relevance of the observed prosody effects in the brain.

## Discussion

The present study aimed to investigate whether and how emotional prosody modulates visual imagery. A new auditorily cued recall task of verbal paired pictures was introduced. At the behavioral level, our results revealed significant prosody effects on the quality of visual imagery regarding accuracy and clarity, and the effects were particularly pronounced in individuals with lower self-reported imagery use tendency. At the neural level, the emotional prosody conditions show stronger activation in various regions including the visual cortex, compared with the neutral condition. Happy vs. frustrated prosody contrast was linked to distinct functional activation in regions including the precuneus and anterior cingulate cortex, which are core components of the default mode network. Interestingly, we found that the prosody-linked activation could predict individual differences in imagery use tendency, further supporting the functional relevance of the prosody effects in the brain. Collectively, these results offer compelling evidence for the modulating effects of emotional prosody on visual imagery and provide new insights into the impact of imagery use tendency on improving imagery robustness in more natural settings.

### Imagery quality is susceptible to emotional prosody

Our experimental results provided compelling evidence for the impact of emotional prosody on the quality of visual mental imagery. At the behavioral level, we found that under the frustrated prosody condition, participants exhibited a reduction in imagery quality, both in terms of clarity and accuracy, compared to the neutral prosody condition.

In the neural data, whole-brain analysis of the fMRI data revealed significant overlap in brain activation between the frustrated vs. neutral and happy vs. neutral conditions. In both contrasts, significant activations were observed in the visual cortex, particularly in the calcarine cortex, and the superior temporal gyrus (STG). Although activation in the inferior frontal gyrus (IFG) was only observed in the frustrated vs. neutral condition, when we used this identified structure as a region of interest, we did observe a stronger BOLD signal in the happy condition compared to the neutral condition (For specific results see Fig. S3).

The visual cortex, particularly the primary visual cortex, has been extensively utilized in previous research for encoding detailed visual information in mental imagery (Albers et al., 2013; Dijkstra et al., 2017; Koenig-Robert & Pearson, 2019; Naselaris et al., 2015). Furthermore, prior research has directly linked the level of activation in the visual cortex to the vividness of mental imagery (Cui et al., 2007). It is reasonable to hypothesize that the observed activation in this region is a result of participants in the emotional prosody condition representing more detailed visual information in their mental imagery.

Nevertheless, we should notice that the heightened signal observed in the visual cortex may not be advantageous. Such activity may be attributed to the demands for the emotional regulation, either resulted from the emotional stimulus itself or the emotional state it triggers. The natural attraction of emotional stimuli to attention has received substantial support in prior research (Pessoa, 2009). It is important to note that under the emotional prosody condition, the representation of this additional information is likely due to weakened suppression of irrelevant information, possibly caused by limited attention. The explanation for this decrease in signal-to-noise ratio is consistent with findings which exhibited a negative correlation between the excitability level of the visual cortex and imagery strength (Rebecca & Pearson, 2015). In other words, the observed decrease in clarity and accuracy in our behavioral results may be attributed to the substantial representation of irrelevant visual information occupying a significant portion of the visual cortex, which is considered to be the “blackboard” of imagery (Albers et al., 2013; Klein et al., 2000; Mechelli, 2004).

Regarding the ROIs that do not significantly contribute to predicting SUIS based on the SHAP results, they are more likely to be primarily involved in emotion-related functions. Previous literature (Morawetz et al., 2016; Underwood et al., 2021) has highlighted the role of the inferior frontal regions in emotion, and considering the discernible emotionality results from the behavioral experiment, the activity in this region could possibly be related to emotional control functions. However, this high-level region may also be associated with other cognitive functions, such as executive control (Rossi et al., 2008), which is expected to play a role in imagery tasks (Pearson, 2019). The precise functions undertaken by the IFG remain a subject for further investigation.

Addressing the role played by STG observed in both the happy vs. neutral and frustrated vs. neutral contrasts, since the analyzed data segments did not include the stage of auditory stimuli perception, it is less likely that the activation in this region is due to differences in the acoustic parameters of the prosody itself. Additionally, given the low contribution of STG to predicted SUIS scores, this activation in the STG is possibly to be related to emotion recognition. As for whether the activation in the STG reflects a weakening of participants’ resistance to cross-modal interference, since prior research has indeed found STG deactivation in normal circumstances during visual imagery tasks as a shielding mechanism (Amedi et al., 2005; Daselaar et al., 2010), this requires further investigation.

Our study provides a potential neural mechanism explanation for how emotional prosody influences the quality of visual imagery. Further research is warranted to gain a more dynamic and systematic understanding of prosody’s effects on imagery.

### DMN facilitates imagination under increased challenges

Our behavioral results indicated that the frustrated condition led to worse imagery quality compared to the neutral condition, whereas the happy condition performed similarly to the neutral condition. To further investigate these specific effect differences related to the type of emotional prosody, we conducted a contrast analysis between the happy and frustrated prosody conditions using the fMRI data. The results revealed significantly positive activations in the left anterior cingulate cortex, right precuneus, and right precentral cortex (see Fig.4(b)). The anterior cingulate cortex and precuneus are crucial components of the Default Mode Network (DMN). Prior research on the neural mechanisms of emotional states has demonstrated that the DMN plays a vital role in distinguishing various emotional states and is primarily involved in the formation of abstract emotional experiences (Satpute & Lindquist, 2019). Therefore, the activation of the DMN may be related to significant differences in emotional experiences. What is even more intriguing, however, is that the DMN also plays a significant role in scene construction during visual imagination tasks (Addis et al., 2007; Daselaar et al., 2010; Hassabis et al., 2007; Hassabis & Maguire, 2007; Zvyagintsev et al., 2013). These findings suggest that compared to the frustrated prosody condition, participants may have been more adept at utilizing the DMN to form emotional experiences under the influence of happy prosody. This, in turn, provides a neural basis for participants to better engage in scene construction, enabling them to “piece together” correct imagery. Interestingly, this inference is further corroborated in subsequent predictive analyses regarding SUIS scores from SHAP model.

However, it should be noted that the brain activity in the anterior cingulate cortex, was not prominent in happy vs. baseline condition (see Fig.5 (c)). This could be attributed to the nature of the baseline condition, where participants fixated their gaze. Even though we instructed participants to empty their minds, they may still have been engaged in episodic memory recall or thoughts related to personal future events, which can activate the DMN (Addis et al., 2007; Daselaar et al., 2010). Hence, the baseline condition itself may have elicited relatively strong DMN activation. Therefore, the DMN regions exhibiting similar or slightly higher activation levels compared to baseline under the happy condition likely indicate that participants effectively utilized the DMN for scene construction. On the other hand, the reduction in DMN activation under the frustrated condition compared to baseline suggests that participants may have struggled to engage this network. Our findings further provide evidence for the role of the DMN in internal scene construction, particularly in situations where visual representations are more complex, and tasks are more challenging.

**Fig. 5.**
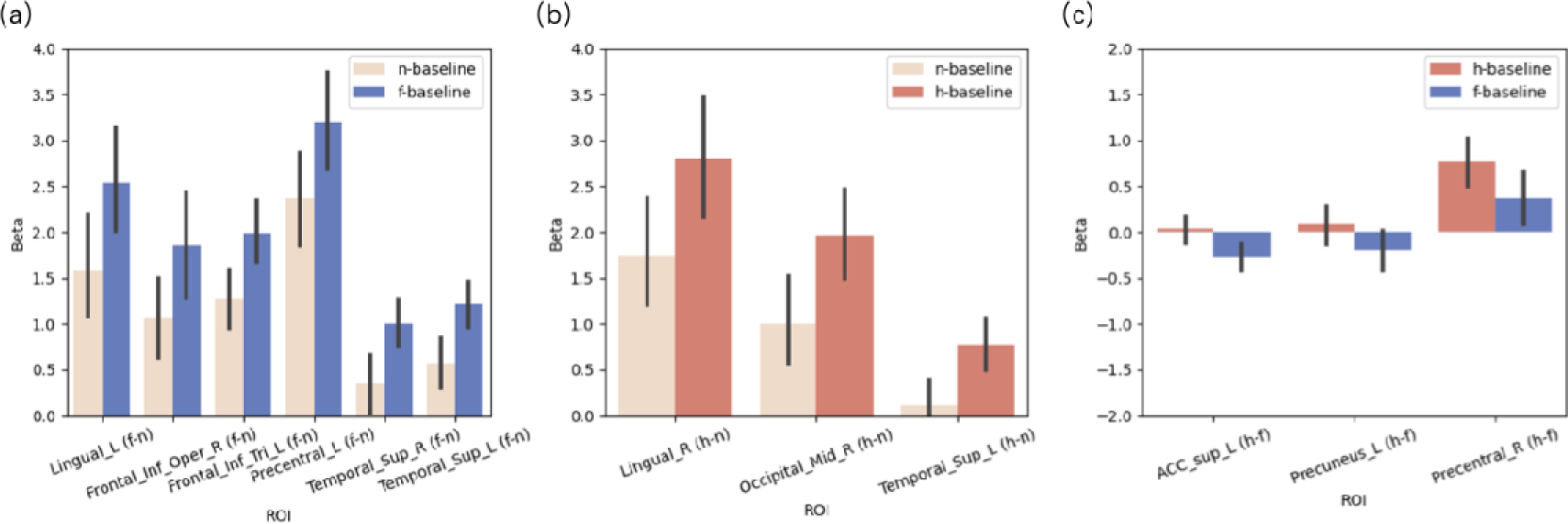
Extracted beta values for ROIs. The ROIs were generated from the f-n, h-n, and h-f contrasts, representing the survived clusters. Beta values were extracted from the corresponding prosody vs. baseline SPM files within the ROIs for each participant. The error bars represent the 95% confidence intervals. (a) Bar plot showing the extracted beta values in the ROIs resulting from the f-n contrast. (b) Bar plot showing the extracted beta values in the ROIs resulting from the h-n contrast. (c) Bar plot showing the extracted beta values in the ROIs resulting from the h-f contrast.

**Fig. 6.**
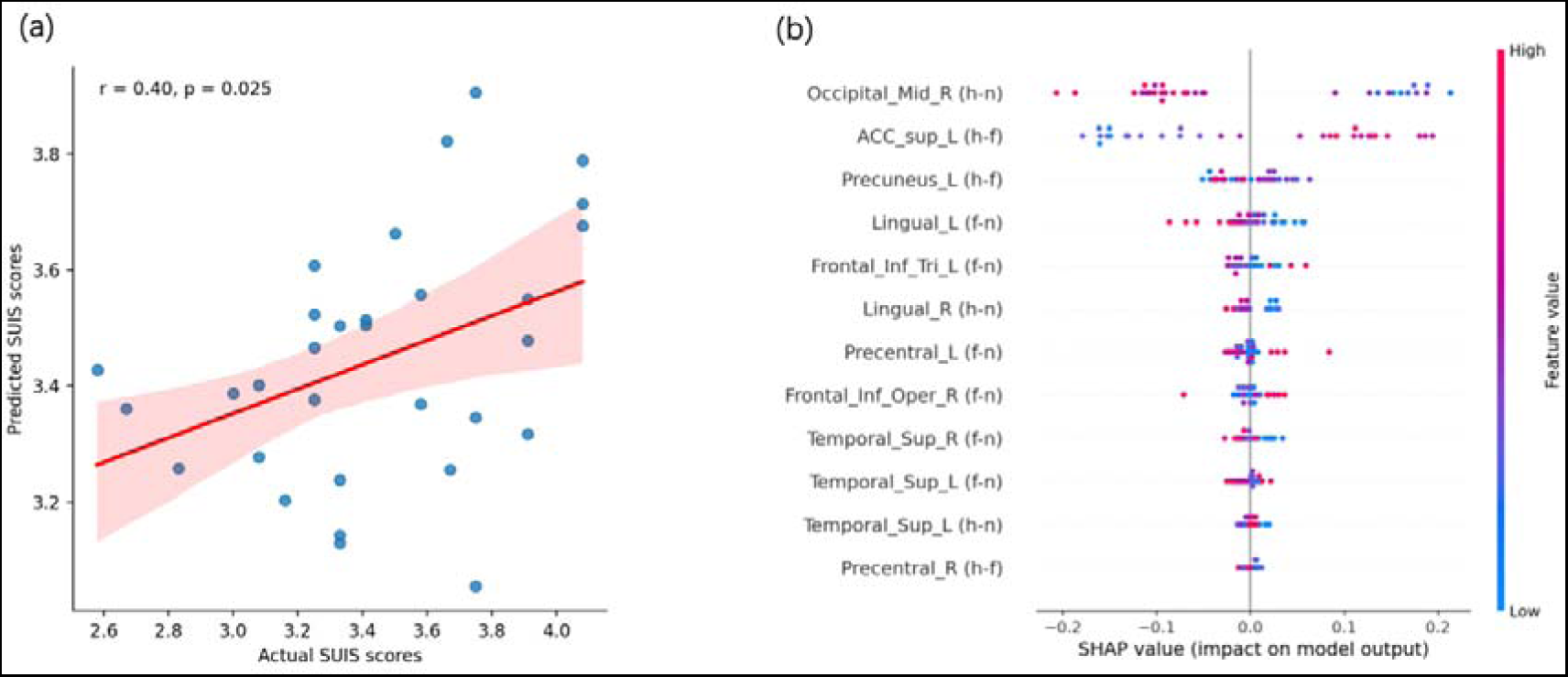
Random Forest Model Performance and SHAP values for each ROI in the model. (a) Scatter plot illustrating the relationship between actual SUIS scores (x-axis) and predicted scores (y-axis) generated by the model. Mean beta values for each ROI were extracted from the SPM files resulting from the corresponding prosody contrasts (f-n, h-n, h-f). These beta values were used to construct a predictive model for SUIS scores using a random forest model with Leave-one-out cross-validation. The plot includes a fitted line with 95% confidence intervals for the scatter points. P-values and correlation coefficients are annotated in the top-left corner of the graph. (b) Scatter plot illustrating the SHAP values of each ROI. The SHAP model was employed to demonstrate the importance of each ROI activity in predicting SUIS scores. Each dot represents a participant’s contribution. The colors at the two ends of the color bar indicate features with larger absolute values.

In addition, we should also note that although both the behavioral emotionality ratings and partial neural brain activation data suggest the presence of emotional activation in participants under the influence of emotional prosody, we did not find activation in the limbic system, especially in the amygdala. This could be due to the fact that limbic regions may not significantly contribute to distinguishing emotional states (Dolan, 2002). Moreover, the absence of amygdala activation could also be attributed to the cognitive demand in our imagery construction task, whose demanding requirement might reduce the amygdala’s response (Erk et al., 2007). Additionally, the amygdala can quickly adapt to emotional stimuli, making it challenging to detect amygdala effects in fMRI data collected after auditory stimuli (Wiethoff et al., 2009).

### Use it or lose it

An intriguing aspect of our results was the influence of individual differences in participants’ imagery use tendency in daily life. At the behavioral level, we observed significant differences in imagery quality between prosody conditions in participants with lower SUIS scores. However, such differences were no longer evident in participants with higher SUIS scores. This suggests that individuals who more frequently use imagery are less susceptible to the influence of emotional stimuli in the environment.

At the neural level, we also found evidence supporting the “use it or lose it” principle in the cognitive ability of imagery. That is, individuals with lower use tendency are more likely to activate irrelevant visual representations in the presence of emotional prosody, and exhibit weaker utilization of the DMN for scene construction according to the SHAP results. This finding also helps us understand why individuals with lower SUIS scores (reflecting imagery use tendency) also tend to have lower Vividness of Visual Imagery Questionnaire (VVIQ) scores (reflecting imagery ability) (Pearson, 2014).

Exploring these individual variations adds new dimensions to existing models of imagery, which may have important implications for understanding emotion-imagery interactions in more manageable approach and in more diverse populations.

## Conclusion

Our findings highlight the importance of considering emotional prosody processing as a critical component in the cognitive theory of mental imagery. Consistent with the “use it or lose it” principle in the biological realm, our results suggest that the more frequently individuals use imagery, the less susceptible their mental imagery is to the impact of such cross-modal emotional stimuli. The incorporation of emotional prosody enables us to simulate more natural scenarios for imagery construction, and sheds light on the mechanism of how our imaginative capabilities adjust to more ecological contexts and how relevant dysfunctions lead to brain disorders.

## Competing interests

The authors indicate no competing interest.

## Supporting information

Supplementary Materials for the main text

## Acknowledgements

Xiang-Zhen Kong was supported by the STI 2030 - Major Project (2021ZD0200409), National Natural Science Foundation of China (32171031), Fundamental Research Funds for the Central Universities (2021XZZX006), and Information Technology Center of Zhejiang University.

## Data availability

The authors will supply the relevant data and code in response to reasonable requests.

## Author contributions

Conceptualization: W.H., X.K.; Methodology: W.H., X.K, Y.P. Experiment design: W.H., Y.P., X.K. Data collection: F.Z., C.Z, C.W., W.H., S.Z. Data Analysis: W.H. Visualization: W.H. Writing-originally draft: W.H. Writing-review & editing: F.Z., C.Z, C.W., W.H., S.Z., Y.P., X.K. Funding Acquisition: Y. P., X.K.

## Notes

### Competing Interest Statement

The authors have declared no competing interest.

