## Supplementary Materials for the main text for "Emotional prosody modulates visual mental imagery"

**Supplemental materials of Huang et al., Emotional prosody modulates visual mental imagery**

Among the participants recruited for our fMRI experiment, imagery phases were self-paced for 5 participants, while the other 26 participants used a fixed duration of 4 seconds for the imagery phase. We conducted separate analyses with only the 26 participants who used the fixed imagery phase duration.

### Methods

Consistent with the analysis for 31 participants, survived clusters under frustrated vs. neutral and happy vs. neutral conditions were generated with uncorrected $p < 0.001$ and passed cluster-level FDR correction at $p < 0.05$. The results closely resemble those obtained from the analysis of 31 participants.

For the statistical analysis under the frustrated vs. happy condition, criteria were set at uncorrected $p < 0.005$, with the additional requirement of passing cluster-level FDR correction at $p < 0.05$. However, no survived clusters were identified in this condition contrast. The absence of significant activation in crucial components of the Default Mode Network (specifically, the anterior cingulate cortex and precuneus) in the results for N=26 may be attributed to the smaller sample size in the frustrated vs. happy condition.

Following the methodology outlined in the main text for training the random forest model, we used three survived clusters from the frustrated vs. neutral condition as masks to extract beta values from the SPM file generated under this contrast. Similarly, we used three survived clusters from the happy vs. neutral condition as masks to extract beta values from the SPM file under this contrast. These beta values were utilized as features to predict SUIS scores for each participant. Unfortunately, we were unable to train a sufficiently effective model, likely due to the small sample size of only 26 participants and the limited number of features (6).

To gain further insights into the relationship between each ROI and SUIS scores, we conducted Pearson correlation analyses for each ROI individually.

### Results

The results revealed patterns remarkably similar to those derived from the analysis of 31 participants (see Tab.S1). Specifically, a significant negative correlation was observed between the visual cortex and SUIS scores in the data from 26 participants. This manifested as a negative correlation in the right Calcarine cortex under frustrated vs. neutral conditions ($p = 0.03$, $r = -0.43$) and in the right middle occipital cortex under happy vs. neutral conditions ($p$=0.04, $r$=-0.4) (see Fig.S4). Notably, these results align with the findings from the dataset of N=31, where SHAP values indicated a significant negative contribution of visual cortex activity to predicted SUIS scores.

**Tab.S1 Significant clusters for prosody condition contrasts.** "n," "f," and "h" represent the neutral, frustrated, and happy prosody conditions, respectively. All clusters were generated using an uncorrected $p$-value threshold of <0.001 and corrected for false discovery rate (FDR) at the cluster level ($p < 0.05$).

| Contrast | Cluster | Cluster size (voxels) | Peak MNI coordinates | *t*-value | Cluster *p*-value (FDR) |
|---|---|---|---|---|---|
| f-n | Calcarine_R | 1147 | 3 -82 2 | 6.05 | 0.000 |
| | Frontal_Inf_Tri_L | 69 | -37 16 22 | 5.06 | 0.003 |
| | Temporal_Sup_R | 120 | 58 -22 -5 | 4.97 | 0.000 |
| h-n | Lingual_R | 480 | 6 -87 -5 | 6.22 | 0.000 |
| | Temporal_Sup_L | 53 | -64 -22 5 | 5.71 | 0.015 |
| | Occipital_Mid_R | 68 | 23 -97 8 | 5.37 | 0.006 |

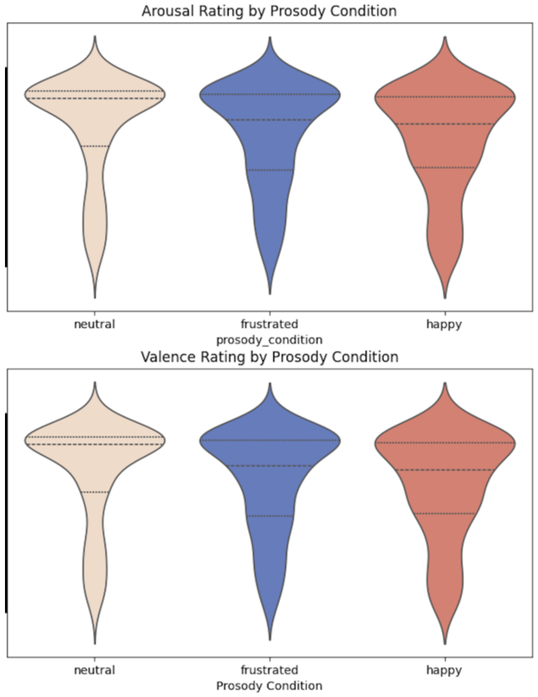


**Fig.S1 Emotionality rating results for all participants from the behavioral experiment (N=39)**, depicting arousal and valence ratings in violin plots. Quartiles are represented by dashed lines. Density estimation is applied beyond the original rating scales. Only trials in which participants reported clear perception were included in the analysis. Wilcoxon non-parametric tests were conducted to examine group differences between prosody conditions, with significant differences denoted by directly labeled p-values.

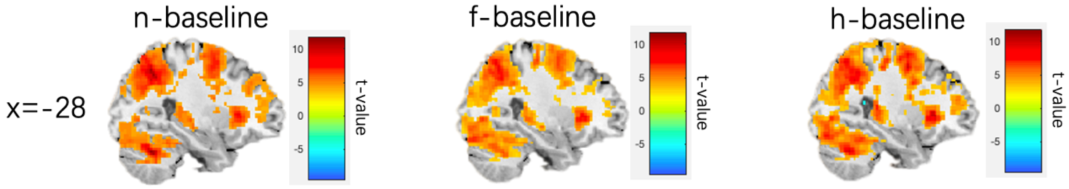


**Fig.S2 Activations observed in each prosody condition compared to baseline (N=31)**. Widespread brain regions exhibited significant activation in these contrasts, including the prefrontal-parietal network, visual cortex, and hippocampus, consistent with previous findings in imagery research.

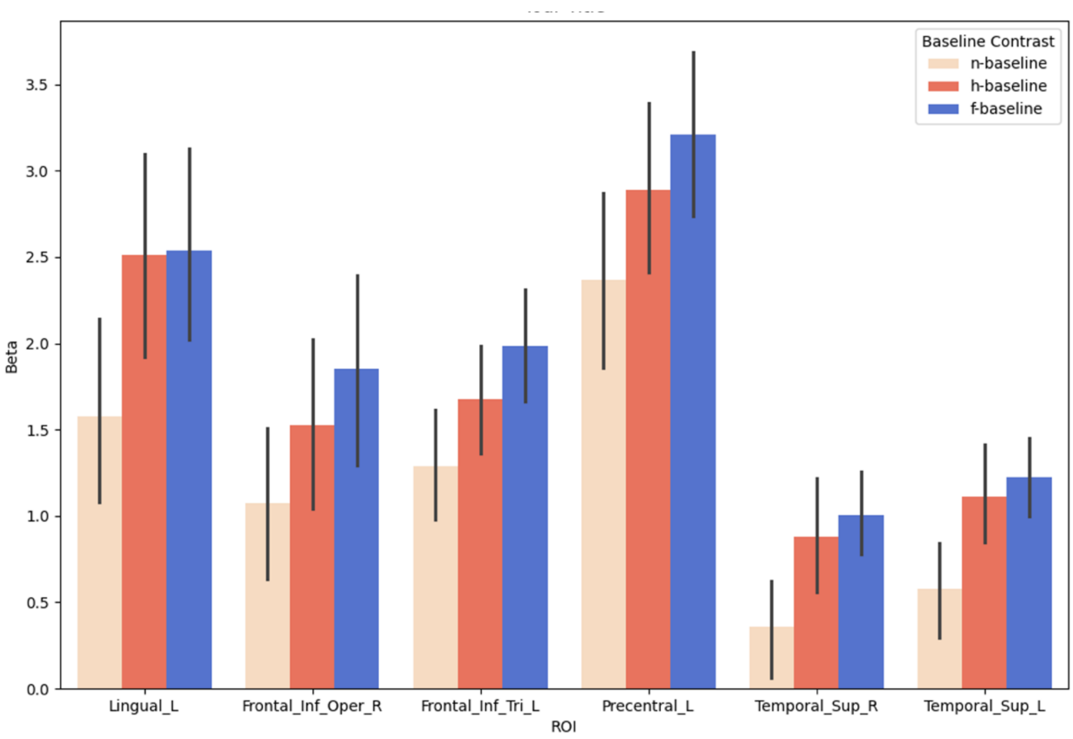


**Fig.S3 Extracted beta values for ROIs.** The ROIs were generated from the frustrated vs. neutral condition. ROIs were generated with uncorrected $p < 0.001$ and all passed cluster-level FDR correction at $p < 0.05$. Beta values were extracted from the corresponding prosody vs. baseline SPM files within the ROIs for each participant. The error bars represent the 95% confidence intervals.

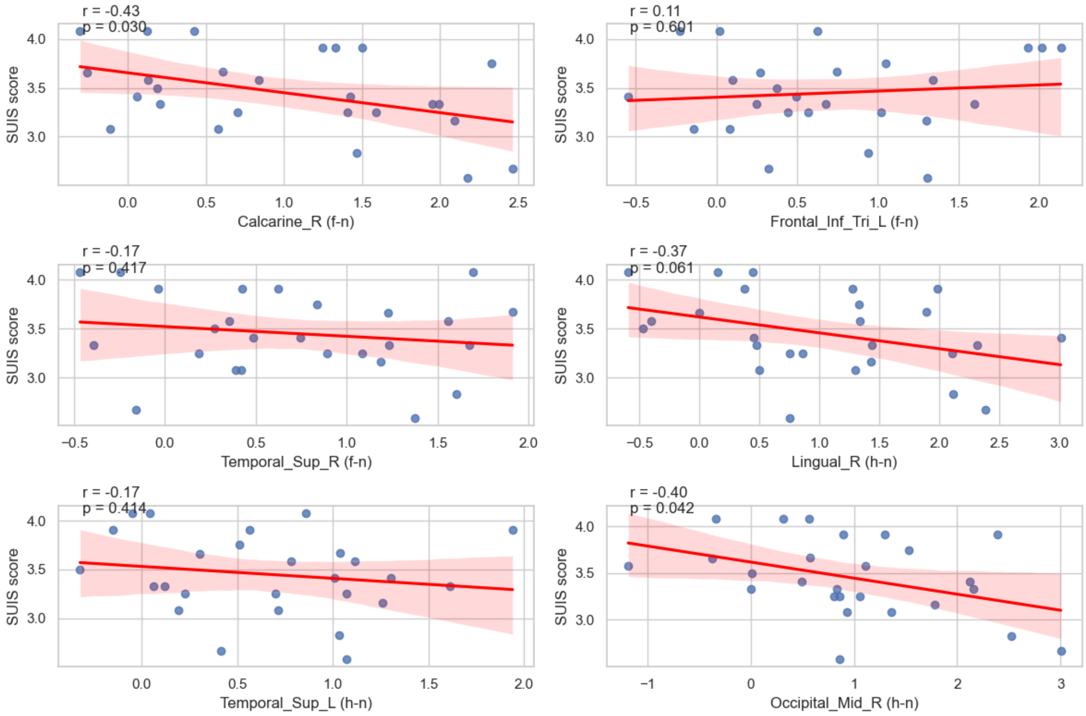


**Fig.S4 Scatter plot illustrating the results of Pearson correlation analysis between beta values for each survived Region of Interest (ROI) and SUIS scores for each participant (N=26)**. The x-axis represents beta values, derived from group-level T-maps comparing frustrated vs. neutral conditions and happy vs. neutral conditions based on 26 participants. The T-maps from frustrated vs. neutral conditions and happy vs. neutral conditions were generated at $p < 0.001$ and passed cluster-level FDR correction at $p < 0.05$. The abbreviations “f,” “h,” and “n” on each subgraph denote frustrated, happy, and neutral conditions, respectively. The plot includes a fitted line with 95% confidence intervals for the scatter points. P-values and correlation coefficients are annotated in the top-left corner of each subgraph.
